# Genome-wide association analysis of cotton salt stress response-related sites

**DOI:** 10.1101/2021.09.01.458514

**Authors:** Juyun Zheng, Zeliang Zhang, Zhaolong Gong, Yajun Liang, Zhiwei Sang, YanChao Xu, Xueyuan Li, Jungduo Wang

**Author notes:** Equal contributors These authors have contributed equally to this work (Juyun Zheng, Zeliang Zhang, Zhaolong Gong).

## Abstract

Soil salinization is the main abiotic stress factor affecting agricultural production worldwide, and salt stress has a significant impact on plant growth and development. Cotton is one of the most salt-tolerant crops. Its salt tolerance varies greatly depending on the variety, growth stage, organs, and soil salt types. Therefore, the selection and utilization of excellent salt-tolerant germplasm resources and the excavation of excellent salt-tolerant salt and salt resistance genes play important roles in improving cotton production in saline-alkali soils. In this study, we analysed the population structure and genetic diversity of 144 elite Gossypium hirsutum cultivar accessions collected from around the world, and especially from China. Illumina Cotton SNP 70K was used to obtain genome-wide single-nucleotide polymorphism (SNP) data for 149 experimental materials, and 18,432 highly consistent SNP loci were obtained by filtering. PCA (principal component analysis) indicated that 149 upland cotton materials could be divided into 2 subgroups, including subgroup 1 with 78 materials and subgroup 2 with 71 materials. Using the obtained SNP and other marker genotype test results, under salt stress, the salt tolerance traits 3d_Germination_potential, 3d_Bud_length_drop_rate, 7d_Germination_rate, 7d_Bud_length_drop_rate, 7d_Germination_weight, 3d_Bud_length, 7d_Bud_length, relative_germination_potential, Relative_germination_rate, 7d_Bud_weight_drop_rate, Salt tolerance index 3d_Germination_potential_index, 3d_Bud_length_index, 7d_Bud_length_index, 7d_Bud_weight_index, and 7d_Germination_rate_index were evaluated by genome association analysis. A total of 27 SNP markers closely related to salt tolerance traits and 15 SNP markers closely related to salt tolerance index were detected. At the SNP locus associated with the traits of the bud length decline rate at 7 days, alleles Gh_A01G0034 and Gh_D01G0028 related to plant salt tolerance were detected, and they are related to intracellular transport, membrane microtubule formation and actin network. This study provides a theoretical basis for the selection and breeding of salt-tolerant upland cotton varieties.

---

Soil salinization is one of the main abiotic stress factors affecting agricultural production worldwide, and salt stress has significant impacts on plant growth and development. Under salt treatment, seed germination, root length, plant height, and fruit development are significantly inhibited (Liang et al, 2014). Therefore, screening and utilizing excellent salt-alkali-tolerant germplasm resources and mining excellent salt-tolerant and salt-tolerant genes play important roles in the transformation and utilization of saline-alkali land and improving the level of agricultural production in saline-alkali land. The damage to cotton caused by salt stress is mainly related to the effects of salt ions on the structure and function of young organs or cell membranes during the developmental transition period, which inhibits the growth of cotton seedlings, affects the growth process, reduces the number of total fruit nodes, and reduces yield and quality (Longenecke, 1973; Longenecke, 1974; Razzouk, et al, 1991).

High-throughput genotyping platforms play an important role in plant genome research. Cai et al. constructed a high-density 80K SNP cotton chip that contained 77,774 SNP sites, among which 352 cotton materials were analysed, and 76.51% of the sites were polymorphic. The chip was utilized to perform a GWAS analysis of 288 upland cotton materials, and total of 54,588 SNPs related to 10 salt tolerance traits were identified, of which 8 SNPs were significantly associated with 3 salt tolerance traits (Cai et al, 2017). Huang et al. used the US 63K chip to perform GWAS analysis on 503 upland cotton materials (63K) and identified 324 SNPs and 160 QTLs related to 16 agronomic traits, of which 38 related areas control 2 or more traits (Huang et al, 2017). Paterson et al. used a population of upland cotton (440 materials) and a population of sea island cotton (219 materials) and the genotyping-by-sequencing (GBS) method to develop 10,129 SNP markers and obtained monomer domains in the whole gene range through analysis. Haplotypes and these results indicate the important role of population genetic methods in the selection of genomic regional variation in the process of cotton domestication (Paterson et al, 2012).

Breeding salt-tolerant crop varieties is the only way to achieve sustainable agricultural development in the future. However, the salt tolerance of plants is a very complicated process. In this study, 124 upland cotton varieties (lines) were used as materials, and 70K SNP chips were used to screen SNP loci and perform genome-wide association analysis on the traits related to salt tolerance at the seedling stage to find significant association sites related to salt tolerance. This study provides a reference and basis for further theoretical studies, such as the isolation of related genes and molecular marker-assisted selection of cotton salt tolerance.

## 1 Materials and methods

### 1.1 Test materials

We sampled 144 modern G. hirsutum cultivars collected from the Chinese national medium-term cotton gene bank at the Institute of Cotton Research (ICR) of the Chinese Academy of Agricultural Sciences (CAAS) (Table 1).

**Table 1.** Sample number and name

| Number Breed |  | Number Breed |  | Number Breed |  |
| --- | --- | --- | --- | --- | --- |
| 1 | Xinluzao NO.1 | 51 | Xinluzhong NO.2 | 101 | Coker310 |
| 2 | Xinluzao NO.2 | 52 | Xinluzhong NO.6 | 102 | Dunhuang 77-116 |
| 3 | Xinluzao NO.3 | 53 | Xinluzhong NO.7 | 103 | Shan 63-1 |
| 4 | Xinluzao NO.4 | 54 | Xinluzhong NO.8 | 104 | Shanmian NO.9 |
| 5 | Xinluzao NO.5 | 55 | Xinluzhong NO.9 | 105 | G164 2 20 |
| 6 | Xinluzao NO.7 | 56 | Xinluzhong NO.10 | 106 | Jimian NO.11 |
| 7 | Xinluzao NO.8 | 57 | Xinluzhong NO.11 | 107 | Lumian NO.17 |
| 8 | Xinluzao NO.9 | 58 | Xinluzhong NO.12 | 108 | Lumian NO.28 |
| 9 | Xinluzao NO.10 | 59 | Xinluzhong NO.14 | 109 | Lu 24 |
| 10 | Xinluzao NO.11 | 60 | Xinluzhong NO.15 | 110 | Lu 25 |
| 11 | Xinluzao NO.12 | 61 | Xinluzhong NO.16 | 111 | Lu 34 |
| 12 | Xinluzao NO.13 | 62 | Xinluzhong NO.17 | 112 | Yumian NO.17 |
| 13 | Xinluzao NO.15 | 63 | Xinluzhong NO.19 | 113 | Baimian NO.1 |
| 14 | Xinluzao NO.16 | 64 | Xinluzhong NO.21 | 114 | Zhong 93001 |
| 15 | Xinluzao NO.17 | 65 | Xinluzhong<br>NO.22 | 115 | CCRI NO.12 |
| 16 | Xinluzao NO.18 | 66 | Xinluzhong<br>NO.26 | 116 | CCRI NO.16 |
| 17 | Xinluzao NO.19 | 67 | Xinluzhong<br>NO.27 | 117 | Zhongmian 41 |
| 18 | Xinluzao NO.20 | 68 | Xinluzhong<br>NO.28 | 118 | CCRI NO.43 |
| 19 | Xinluzao NO.21 | 69 | Xinluzhong<br>NO.30 | 119 | Emian NO.10 |
| 20 | Xinluzao NO.23 | 70 | Xinluzhong<br>NO.32 | 120 | Emian NO.12 |
| 21 | Xinluzao NO.24 | 71 | Xinluzhong<br>NO.34 | 121 | Emian NO.21 |
| 22 | Xinluzao NO.25 | 72 | Xinluzhong<br>NO.35 | 122 | Wanmian 8407 |
| 23 | Xinluzao NO.26 | 73 | Xinluzhong<br>NO.36 | 123 | Sumian NO.8 |
| 24 | Xinluzao NO.27 | 74 | Xinluzhong<br>NO.38 | 124 | Sumian NO.12 |
| 25 | Xinluzao NO.29 | 75 | Xinluzhong<br>NO.39 | 125 | Suyuan 04-129 |
| 26 | Xinluzao NO.30 | 76 | Xinluzhong<br>NO.40 | 126 | Ganmian NO.10 |
| 27 | Xinluzao NO.31 | 77 | Xinluzhong<br>NO.41 | 127 | Ganmian NO.17 |
| 28 | Xinluzao NO.32 | 78 | Xinluzhong<br>NO.42 | 128 | Chuan 7327 20 |
| 29 | Xinluzao NO.33 | 79 | Xinluzhong<br>NO.44 | 129 | Yumian NO.1 |
| 30 | Xinluzao NO.34 | 80 | Xinluzhong<br>NO.45 | 130 | Liaomian NO.9 |
| 31 | Xinluzao NO.35 | 81 | Xinluzhong<br>NO.46 | 131 | Liaomian NO.16 |
| 32 | Xinluzao NO.36 | 82 | Xinluzhong<br>NO.50 | 132 | Dai-80 |
| 33 | Xinluzao NO.37 | 83 | Xinluzhong<br>NO.54 | 133 | Montenegro cotton NO.1 |
| 34 | Xinluzao NO.38 | 84 | Xinluzhong<br>NO.56 | 134 | Pidcotton |
| 35 | Xinluzao NO.39 | 85 | Xinluzhong<br>NO.58 | 135 | MacNair 210 |
| 36 | Xinluzao NO.40 | 86 | Xinluzhong<br>NO.59 | 136 | Huazhong 106 |
| 37 | Xinluzao NO.41 | 87 | Xinluzhong<br>NO.60 | 137 | Keyuan NO.1 |
| 38 | Xinluzao NO.42 | 88 | Xinluzhong<br>NO.61 | 138 | Bu 3363 |
| 39 | Xinluzao NO.45 | 89 | Xinluzhong<br>NO.62 | 139 | Bamian NO.1 |
| 40 | Xinluzao NO.46 | 90 | Xinluzhong<br>NO.63 | 140 | Chad NO.3 |
| 41 | Xinluzao NO.47 | 91 | Xinluzhong<br>NO.64 | 141 | Turkmen upland cotton |
| 42 | Xinluzao NO.48 | 92 | Xinluzhong<br>NO.65 | 142 | Phenphenol phenol NO.1 |
| 43 | Xinluzao NO.49 | 93 | Xinluzhong<br>NO.68 | 143 | Miscot7803-52 |
| 44 | Xinluzao NO.50 | 94 | Xinluzhong<br>NO.69 | 144 | Si-6524 |
| 45 | Xinluzao NO.51 | 95 | Xinlu 201 | 145 | Sparculent H10 |
| 46 | Xinluzao NO.52 | 96 | Xinlu 202 | 146 | Yinmian NO.1 |
| 47 | Xinluzao NO.53 | 97 | Nongken<br>NO.5 | 147 | America 28114-313 |
| 48 | Xinluzao NO.60 | 98 | Shache Soil<br>cotton | 148 | Columbia |
| 49 | Xinluzao NO.61 | 99 | Kuche<br>T94-4 | 149 | Miscot 8711ne |
| 50 | Xinluzhong NO.1 | 100 | Bazhou<br>6510 |  |  |

### 1.2 DNA extraction and genotyping

All cotton seeds were grown in a soil mixture in a fully automated greenhouse under a 12-h light/12-h dark cycle at 28 °C. Total genomic DNA was extracted from 5-day-old seedlings germinated from five well-developed seeds of each accession using a Qiagen DNeasy plant mini kit (Qiagen, CA, USA) following the protocol provided by the manufacturer. Genotyping was conducted at the CapitalBio Technology Platform in China using the Illumina Cotton SNP 70k Beadchip (Illumina, Inc. San Diego, CA 92122 USA). All genotype calls were extracted from the raw data using GenomeStudio (Illumina). Loci with a distance between two significant association sites of less than 10 cm (Wang et al., 2014) were classified as QTLs. The reference genome was:

Gossypium hirsutum (AD1) ‘TM-1’ genome NAU-NBI_v1.1_a1.1

### 1.3 Molecular genetic diversity and phylogenetic analyses

PHYLIP (http://evolution.genetics.washington.edu/phylip.html) was used to calculate the genetic distance matrix of the sample, Notepad++ software was used to adjust the genetic distance matrix file into a suitable format, and the neighbour-joining method was used to construct a phylogenetic tree. After generating the tree file, iTOL (https://itol.embl.de/) was used to draw the evolutionary tree diagram. Principal component analysis (PCA) was performed on cotton population materials using GCTA software (Yang et al, 2011) using the detected SNPs. Then, R software was used to calculate the vector of each principal component and draw the PCA scatter plot.

### 1.4 Population structure and kinship analysis

Population structure was examined using the Bayesian model-based approach implemented in STRUCTURE V2.3.4 (Pritchard et al. 2000). SPAGeDi (Hardy et al, 2002) software was used to estimate the relative kinship between two individuals in a natural population. The kinship itself is the relative value that defines the genetic similarity between two specific materials and the genetic similarity between any material. Therefore, when the kinship value between the two materials is less than 0, it is directly defined as 0.

### 1.5 Linkage disequilibrium analysis

On the same chromosome, the linkage disequilibrium between two SNPs within a certain distance can be calculated (such as 1,000 kb), and the linkage disequilibrium strength is represented by r2. The closer r2 is to 1, the stronger the strength of linkage disequilibrium. The SNP spacing is fit to r2, and a graph can be drawn to represent the variation of r2 with distance. Generally, the closer the SNP spacing is, the larger r2 is, and the farther the SNP spacing is, the smaller r2 is. The distance travelled when the maximum r2 value drops to half is used as the LD decay distance (LDD) of linkage disequilibrium. The longer the LDD is, the smaller the probability of recombination within the same physical distance; the shorter the LDD is, the greater the probability of recombination within the same physical distance. Plink2 (Purcell et al., 2007) software was used for LD analysis.

### 1.6 Association analysis of salt tolerance traits

Linkage disequilibrium analysis of natural populations was used to evaluate traits. Through a certain amount of population SNP marker data, combined with population structure and target trait phenotype data, the target region or site associated with the target trait can be located.

Salt stress conditions and salt-tolerant trait collection: The salt tolerance test during the germination period used double-layer filter paper rolls to stand the plant upright. Two pieces of filter paper each 20 cm in length and width were cut, and one piece of filter paper was spread on the test bench with a sprayer containing NaCl solution. The filter paper was soaked, and 15 seeds were placed 2 cm down from the top of the filter paper. The filter paper was then placed vertically into the culture box. Approximately 30 rolled filter papers were placed in each culture box. The culture box was then placed at 28°C, and the photoperiod was 10h/14h (L/D), with heat preservation and culture in a constant temperature light incubator. The germination potential of seeds and the length of each seed were measured on the 3rd day, and the germination rate, bud length and stem fresh weight of the seeds were measured on the 7th day. This process was repeated 3 times. The treatment concentrations of NaCl solution were 0 NaCl (CK) and 150.00 mmol/L NaCl.

The calculation formula analyses the relative values of the salt stress environment and the control conversion.

Relative germination potential% = germination potential of treated seeds/germination potential of control seeds × 100%

Relative germination rate% = germination rate of treated seeds/germination rate of control seeds × 100%

Decrease rate% = (treatment traits-control traits)/control traits × 100%

Salt tolerance index:

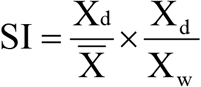

Note: X_d_ and X_w_ are the measured values of a certain index of each material under salt stress conditions and control conditions, respectively, and 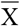 is the average value of this index under salt stress conditions.

SAS software was used to perform the best linear unbiased prediction (BLUP) for salt tolerance traits, TASSEL v5.0 software was used to perform correlation analysis for each trait based on the four models of glm, mlm, cmlm, and fastlmm, and the result of the structure was used as a fixed effect. Among them, the mixed linear model formula of TASSEL software is as follows:

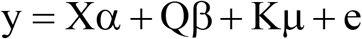

Note: SPAGeDi (Hardy et al., 2002) software was used to calculate the genetic relationship K between samples. The general linear model uses Q population structure information, while the mixed linear model uses Q+K, which is the population structure and genetic relationship information. X is the genotype and Y is the phenotype. In the end, an association result can be obtained for each SNP site.

Salt stress cotton transcriptional group data download: Sequencing of allotetraploid cotton (Gossypium hirsutum L. acc. TM-1) provides a resource for fibre improvement. Nat Biotechnol, 2015, doi: 10.1038/nbt.3207.

### 1.7 Prediction and functional annotation of salt-tolerant candidate genes

The independent significant SNP sites selected by the GWAS analysis results and LD calculations, plus or minus 500 kb upstream and downstream of the physical location of each SNP site as the candidate gene physical location query area were identified by mapping the gene or Arabidopsis homologous gene and annotating information to narrow down the target candidate genes. NCBI, COTTONGEN, CNKI, Tair3 and other websites were used to annotate gene functions and compare homologous sequences.

## 2 Results and analysis

### 2.1 Genetic diversity analysis and population structure

#### 2.1.1 Group structure

After 63,058 probe sequences were blast aligned with the genome, the optimal result screened out was the position of the SNP on the reference genome. The SNP data of 149 experimental materials were detected by Illumina Cotton SNP 70K and filtered according to the minor allele frequency (MAF: 0.05) and site integrity (INT: 0.1), and 18,432 highly consistent SNP sites were obtained. PCA found that 149 individuals of upland cotton could be divided into 2 subgroups, including subgroup 1 (marked in red), consisting of 78 materials, and subgroup 2, consisting of 71 individuals, PC1 9.25% and PC2 5.2%. Based on the analysis of the phylogenetic tree constructed from the SNP data, the 149 materials could be divided into two subgroups, and there was gene exchange between the two subgroup materials. The results are consistent with the grouping structure of PCA (Figure 1). The hybrid model was used in Structure 2.3.4 software. First, the number of subgroups (K) was set to 2–10, and each K value was set to 3 repetitions. Assuming that each site is independent, the Markov chain Monte Cardo (MCMC) at the beginning of the noncount iteration (length of burn in period) was set to 10,000 times, and then the MCMC after no-count iterations was set to 1,000,000 times. The optimal K value was selected according to the principle of maximum likelihood value to determine the number of subgroups and the group structure. In this experiment, using the Q value calculation and Structure software, the population was divided into 2 subgroups.

**Figure 1.**
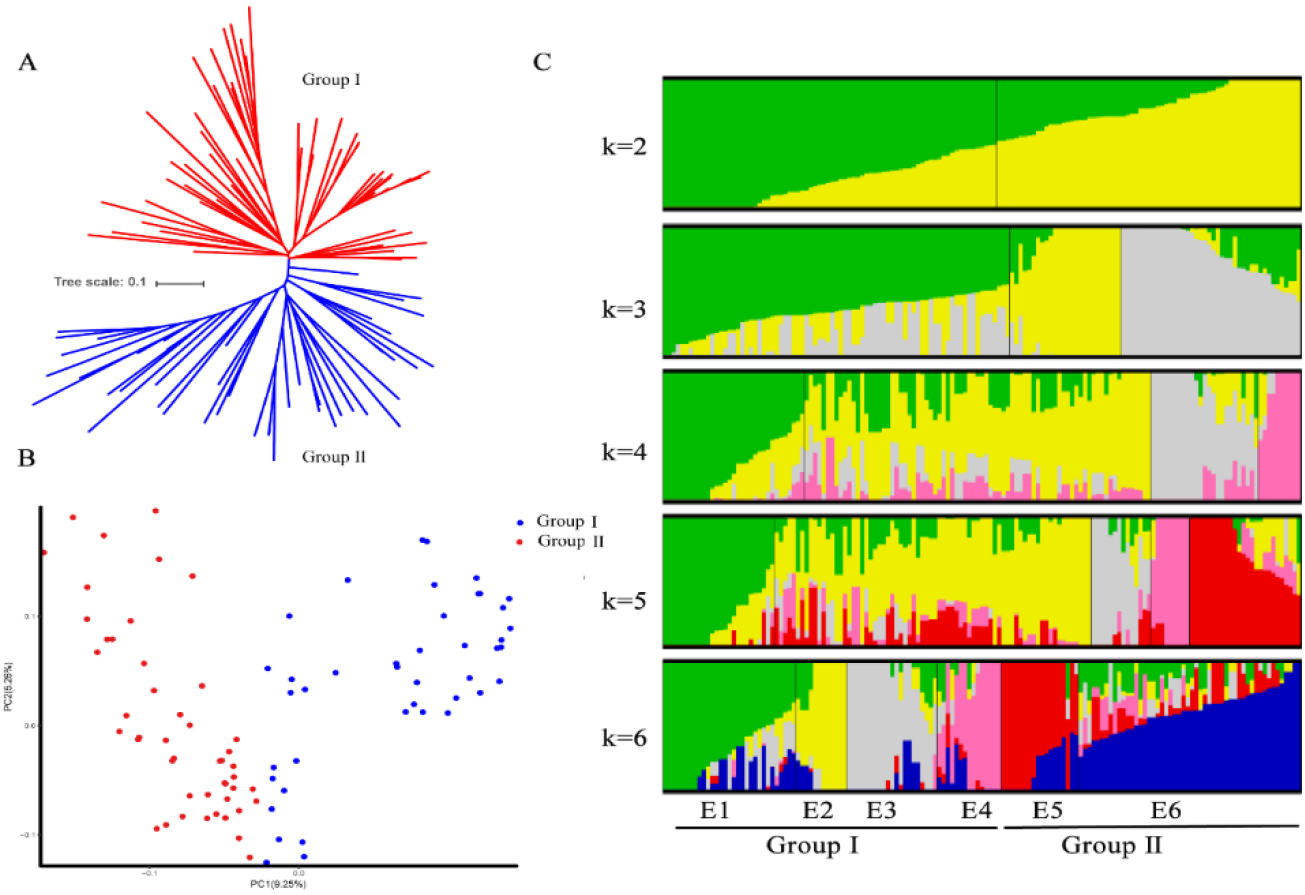
Analysis of group structure. A: Construction of a phylogenetic tree based on SNP typing data. B: PCA; C: structure analysis.

#### 2.1.2 Material heterozygosity

The individual heterozygosity analysis found that 95% of the materials were less than 30% heterozygous, and 80% of the individual materials were less than 5% heterozygous (Figure 2).

**Figure 2.**
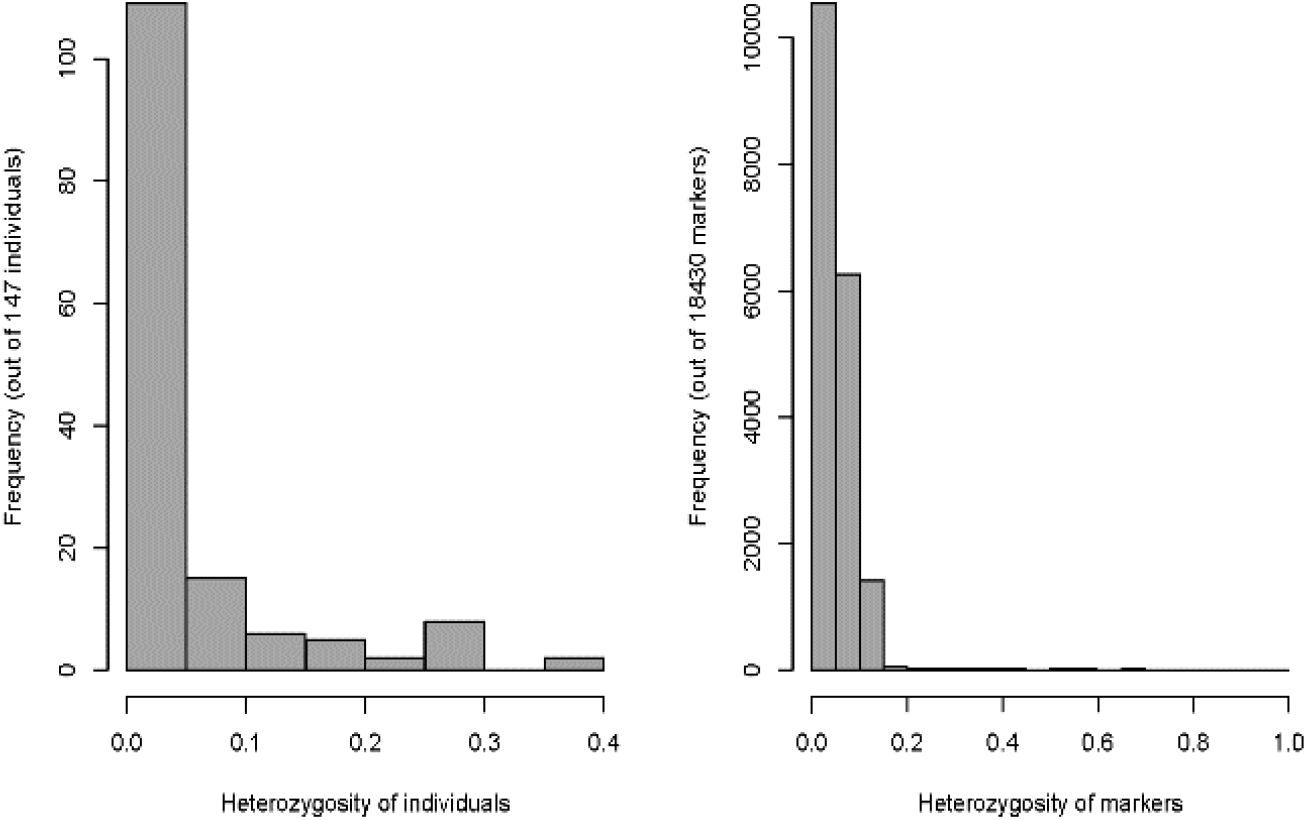
Frequency distribution of material heterozygosity and distribution of labelled heterozygosity

#### 2.1.3 Kinship distribution

Figure 3 shows that the 149 cotton varieties can be divided into 2 subgroups. Among the 149 varieties, the genetic relationships between most varieties were weak (the yellow parts in the figure), and the genetic relationships between a few materials were very close (the dark red parts in the picture).

**Figure 3.**
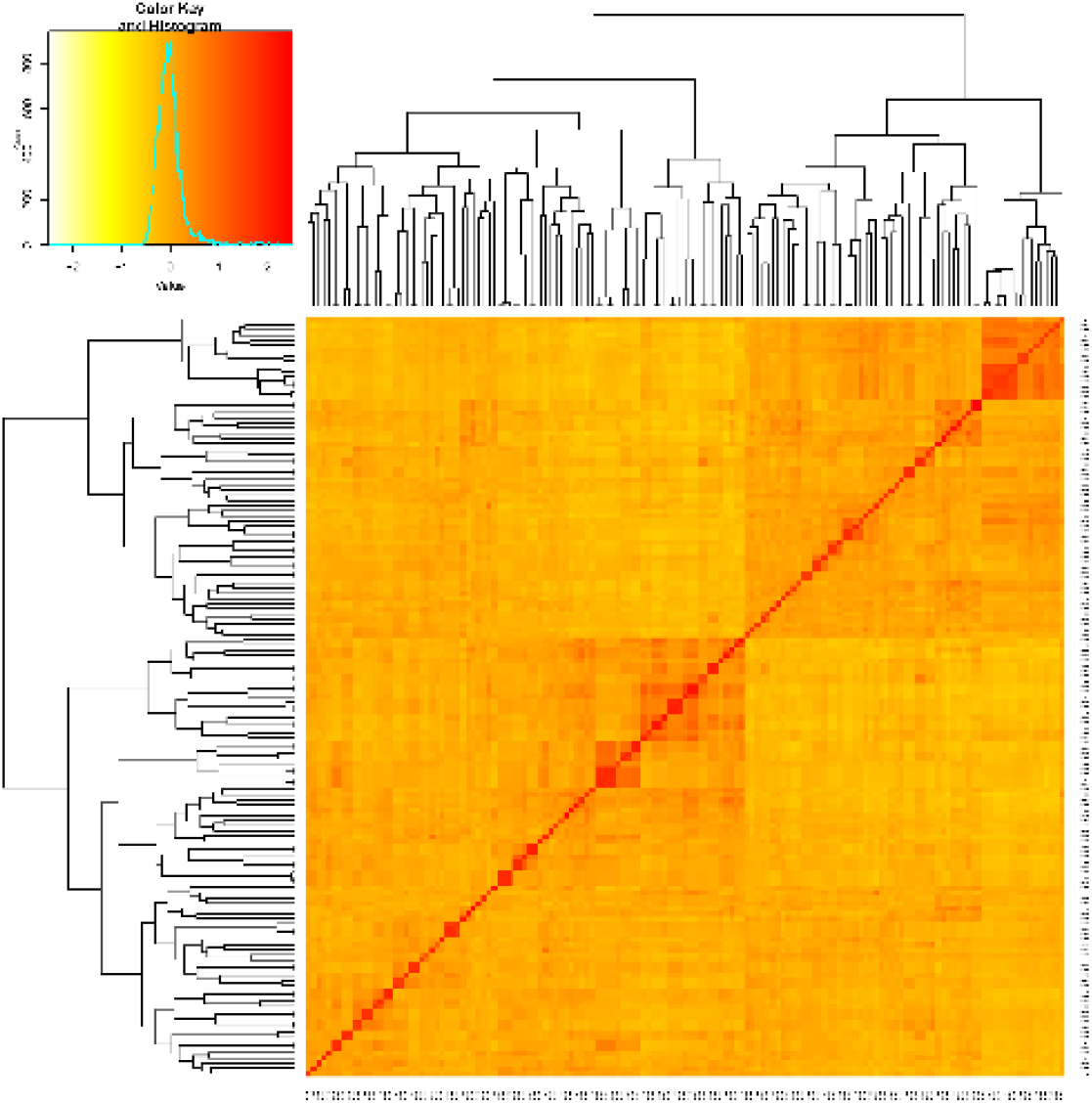
Heat map and cluster analysis based on the genetic relationship of 149 upland cotton varieties

#### 2.1.4 Analysis of linkage disequilibrium

The LD distance decreases as the physical position of the SNP on the chromosome increases. The analysis found that the LD distance of 149 samples was 432 K (R square=0.5) (Figure 4). Slightly higher than the previous study (Wang, 2019), Chinese upland cotton material was 296 kb. This result further shows that the genetic diversity of the selected material is reduced.

**Figure 4.**
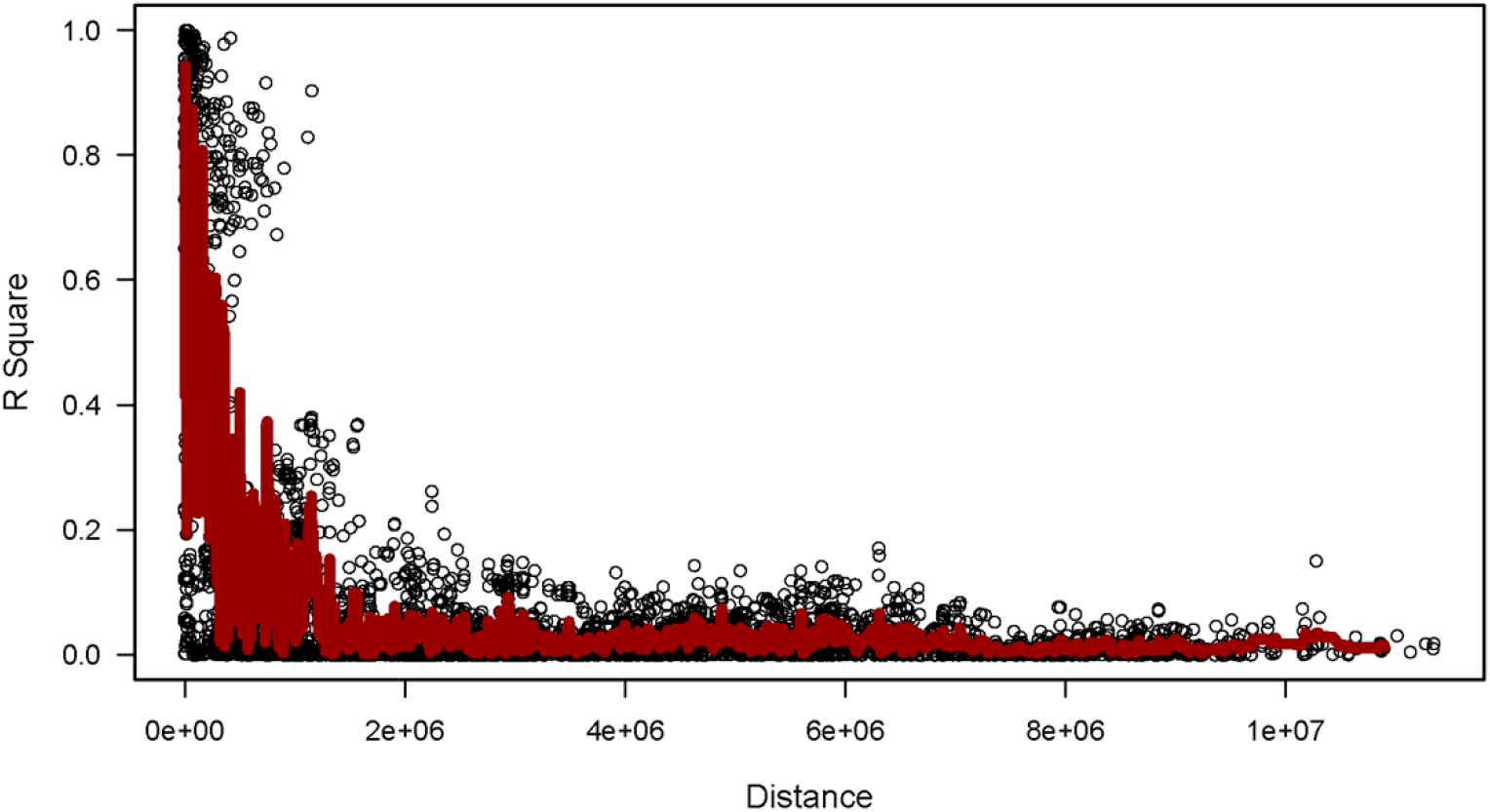
LD attenuation analysis of 144 materials

### 2.2 Phenotypic statistical analysis

In the best linear unbiased prediction (BLUP) for salt tolerance index traits, a total of 5 phenotypic traits in the salt tolerance index, 3d germination vigour index under salt stress, 3d bud length index, 7d bud length index, 7d bud weight index, 7d germination rate index, were identified. Figure 5 shows that the phenotypic distributions of the 5 traits were all normally distributed, indicating that these traits are all typical quantitative traits and are controlled by minor-effect polygenes. Using R language to calculate the Pearson correlation coefficients between traits, it was found that the correlations between different traits were low (Table 3), which may be because these 5 traits are controlled by independent inherited genetic sites in response to salt stress, indicating consistent complexity.

**Figure 5.**
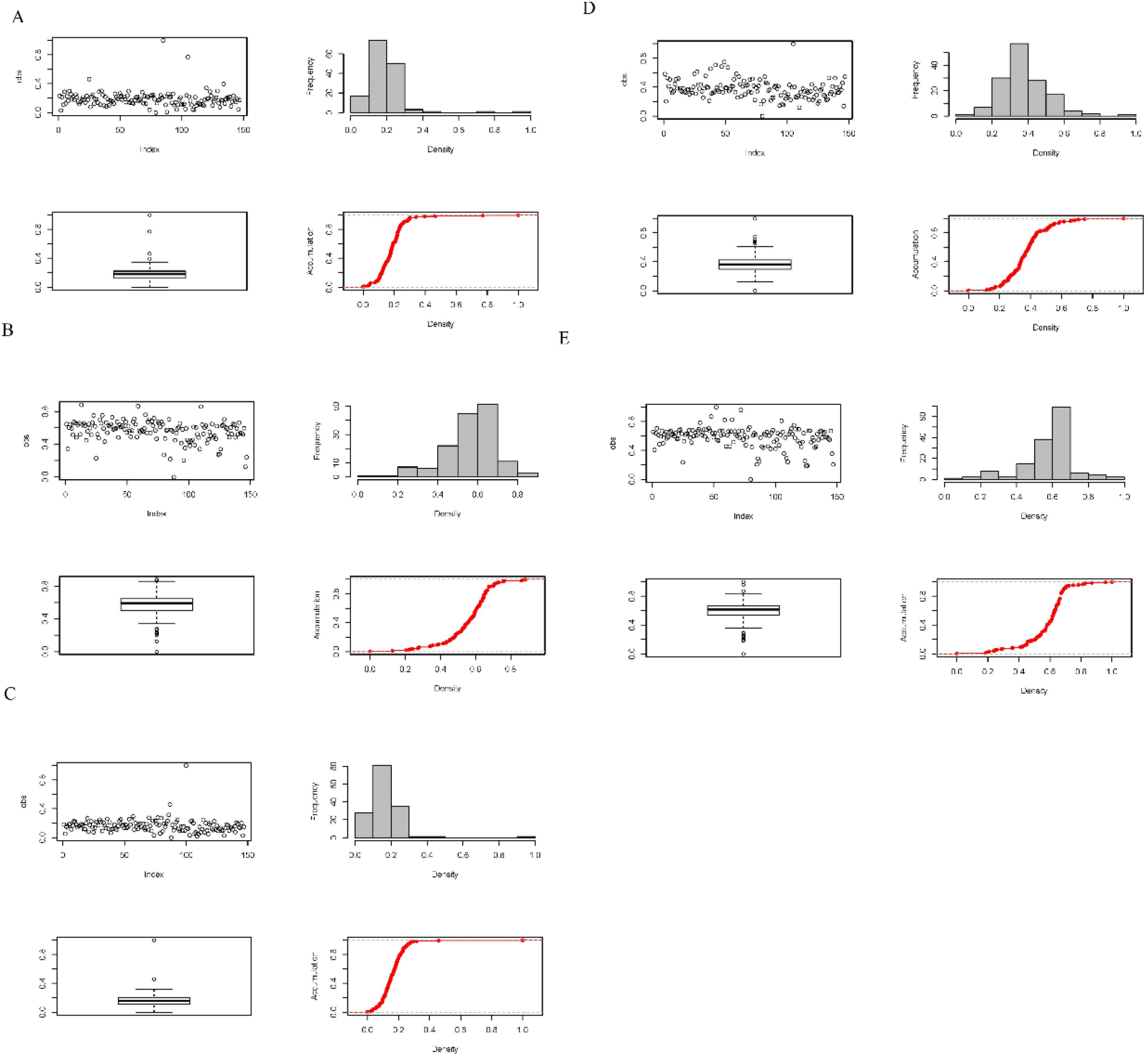
Distribution of traits in the three environments

**Table 2.** Salt tolerance traits and salt tolerance index

|  |  |
| --- | --- |
| 3d_Germination_potential | 3d_Bud_length_drop_rate |
| 7d_Germination_rate | 7d_Bud_length_drop_rate |
| 7d_Germination_weight | 3d_Germination_potential_index |
| 3d_Bud_length | 7d_Germination_rate_index |
| 7d_Bud_length | 7d_Bud_weight_index |
| Relative_germination_potential | 3d_Bud_length_index |
| Relative_germination_rate | 7d_Bud_length_index |
| 7d_Bud_weight_drop_rate |  |

**Table 3.** Correlation analysis of salt tolerance index traits

|  | 3d_Germination_<br>potential_index | 7d_Germinat<br>ion_rate_inde<br>x | 7d_Bud_w<br>eight_inde<br>x | 3d_Bud_l<br>ength_ind<br>ex | 7d_Bud_l<br>ength_ind<br>ex |
| --- | --- | --- | --- | --- | --- |
| 3d_Germinati<br>on_potential_<br>index | 1 | 0.565096 | 0.330095 | 0.006608 | 0.178595 |
| 7d_Germinati<br>on_rate_index | 0.565096 | 1 | 0.484635 | 0.000286 | -0.03288 |
| 7d_Bud_weig<br>ht_index | 0.330095 | 0.484635 | 1 | 0.475708 | 0.192883 |
| 3d_Bud_length_index | 0.006608 | 0.000286 | 0.475708 | 1 | 0.230305 |
| X7d_Bud_length_index | 0.178595 | -0.03288 | 0.192883 | 0.230305 | 1 |

### 2.3 Association analysis of salt tolerance traits

Based on the identification results of the morphological, physiological, biochemical, and yield traits of the specific germplasms of upland cotton under saline-alkali stress, an association analysis of the salt tolerance traits was carried out, and the favourable alleles related to the salt tolerance of the specific germplasm of upland cotton were located. A total of 27 SNP sites related to salt tolerance traits were detected (see Tables 2–12 for details): 3 sites related to 7d_Bud_length; 10 sites related to 7d_Bud_length_drop_rate; sites related to 7d_Germination_rate 3; 3 sites related to 7d_Germination_weight; and 8 sites related to Relative_germination_rate. The marker loci were scattered on 6 cotton chromosomes, A01, D01, D05, D08, D11, and D13, without clustering, and 6 QTLs related to salt tolerance traits were located on different chromosomes (Table 4). The research and development of SNP markers and QTL sites closely related to salt tolerance traits can be applied to the molecular marker-assisted selection of cotton salt tolerance.

**Table 4.** SNP sites related to salt tolerance traits

| Number | Traits | Chromosomes | Position | P value | Threshold |
| --- | --- | --- | --- | --- | --- |
| 1 | 7d_Bud_length | D08 | 64320143 | 1.85E-05 | 4.731944 |
| 2 |  | D13 | 35877478 | 4.63E-05 | 4.334531 |
| 3 |  | scaffold33030 | 93 | 9.89E-05 | 4.004747 |
| 4 | 7d_Bud_length_drop_rate | D01 | 16458717 | 2.58E-07 | 6.588126 |
| 5 |  | A01 | 22140802 | 7.37E-07 | 6.1328 |
| 6 |  | A01 | 22174537 | 7.37E-07 | 6.1328 |
| 7 |  | A01 | 22350123 | 7.37E-07 | 6.1328 |
| 8 |  | A01 | 22376230 | 7.37E-07 | 6.1328 |
| 9 |  | A01 | 22387777 | 7.37E-07 | 6.1328 |
| 10 |  | D01 | 16458204 | 7.37E-07 | 6.1328 |
| 11 |  | D05 | 58225028 | 5.94E-06 | 5.226497 |
| 12 |  | D05 | 58227014 | 1.45E-05 | 4.83936 |
| 13 |  | D11 | 58447888 | 1.73E-05 | 4.762019 |
| 14 | 7d_Germination_r<br>ate | D05 | 12154352 | 2.75E-05 | 4.560614 |
| 15 |  | D05 | 12155655 | 3.04E-05 | 4.516439 |
| 16 |  | A05 | 12181224 | 7.78E-05 | 4.109102 |
| 17 | 7d_Germination_<br>weight | D08 | 2191589 | 5.44E-05 | 4.264409 |
| 18 |  | D05 | 12154352 | 5.55E-05 | 4.255444 |
| 19 |  | D05 | 12155655 | 9.21E-05 | 4.035935 |
| 20 | Relative_germinat<br>ion_rate | D08 | 43495093 | 2.24E-05 | 4.65032 |
| 21 |  | D08 | 43541365 | 2.65E-05 | 4.57704 |
| 22 |  | D08 | 43557843 | 2.91E-05 | 4.5365 |
| 23 |  | D08 | 43483056 | 3.31E-05 | 4.48071 |
| 24 |  | D08 | 43501088 | 3.36E-05 | 4.473257 |
| 25 |  | D08 | 43479511 | 3.48E-05 | 4.458785 |
| 26 |  | D01 | 54316248 | 5.13E-05 | 4.28993 |
| 27 |  | D01 | 54289162 | 7.82E-05 | 4.106987 |

### 2.4 Association analysis of salt tolerance index traits

The results of the GWAS under the optimal model of the salt tolerance index traits under BLUP were counted and explained, and the results are shown in Table 2. A total of 15 significant SNP-trait associations were detected (Table 5). It was also found that among these 5 traits, only 4 traits had significant SNPs, while the 7d_Bud_wight_index did not have a significant locus. This may be because this trait is more complicated and controlled by multiple minor QTLs (Figure 6).

**Table 5.** SNP sites related to salt tolerance index traits

| Traits | SNP | Chromosomes | Position | P value |
| --- | --- | --- | --- | --- |
| 7d_Bud_length_index | rs__A11__85527572 | 11 | 85,527,572 | 5.84E-05 |
| 7d_Germination_rate_index | rs__A10__79275413 | 10 | 79,275,413 | 0.000229 |
|  | rs__D08__43479511 | 21 | 43,479,511 | 0.000485 |
|  | rs__D08__43483056 | 21 | 43,483,056 | 0.000485 |
|  | rs__D08__43495093 | 21 | 43,495,093 | 0.00034 |
|  | rs__D08__43501088 | 21 | 43,501,088 | 0.000485 |
|  | rs__D08__43541365 | 21 | 43,541,365 | 0.000384 |
|  | rs__D08__43557843 | 21 | 43,557,843 | 0.000495 |
| 3d_Germination_potential_index | rs__A05__12472578 | 5 | 12,472,578 | 0.000299 |
|  | rs__A05__12473346 | 5 | 12,473,346 | 0.000675 |
|  | rs__D02__56360435 | 15 | 56,360,435 | 0.000872 |
|  | rs__D05__3861440 | 18 | 3,861,440 | 0.000815 |
|  | rs__D05__3870101 | 18 | 3,870,101 | 0.000887 |
|  | rs__D05__3873693 | 18 | 3,873,693 | 0.000301 |
| 3d_Bud_length_index | rs__D01__883613 | 14 | 883,613 | 0.000872 |

**Figure 6.**
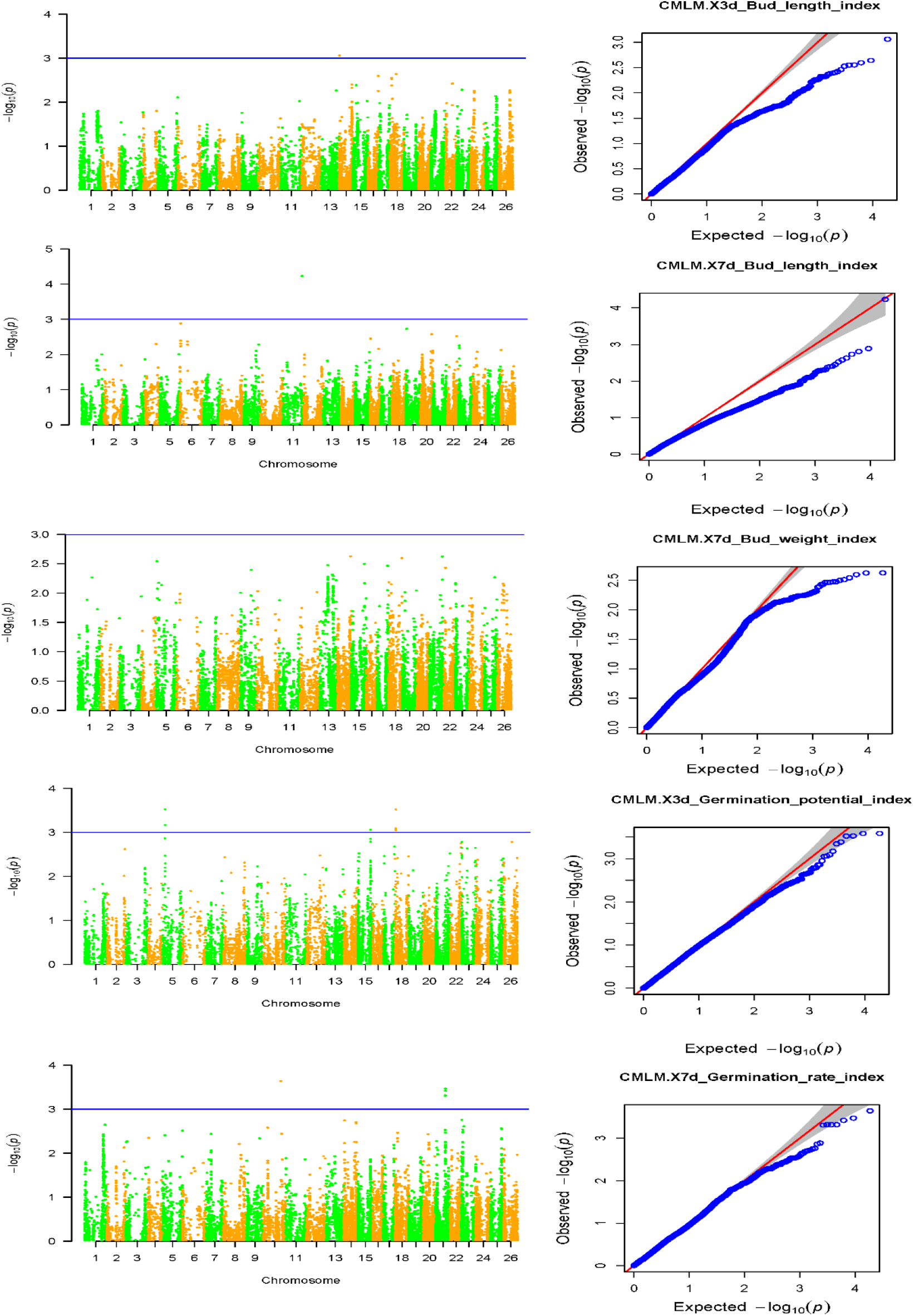
Manhattan and QQ charts of the 5 salt tolerance index traits Note: Manhattan graph: the abscissa represents the position of the chromosome, the ordinate represents the p value (-log10(p)) taking the negative logarithm based on 10, and the scattered dots (or lines) on the graph represent the corresponding data for each SNP site. log10(p). The blue horizontal line is the threshold line. Scattered points (or lines) that exceed the threshold line are candidate sites. QQ chart: The abscissa represents the expected value, and the ordinate represents the observed value. The red line in the figure represents the 45° centre line, and the grey area is the 95% confidence interval of the scattered points in the figure.

### 2.5 Candidate gene screening

Five SNP sites on chromosome A01 and 2 SNP sites on chromosome D01 associated with a decrease in bud length at 7 days were detected. The alleles Gh_A01G0034 and Gh_D01G0028 related to salt tolerance were detected and are located in A01 and D01 by homology. On the chromosome, Gh_A01G0034 and Gh_D01G0028 all reached their peak transcriptome expression at 12 h of salt stress (Figure 7).

**Figure 7.**
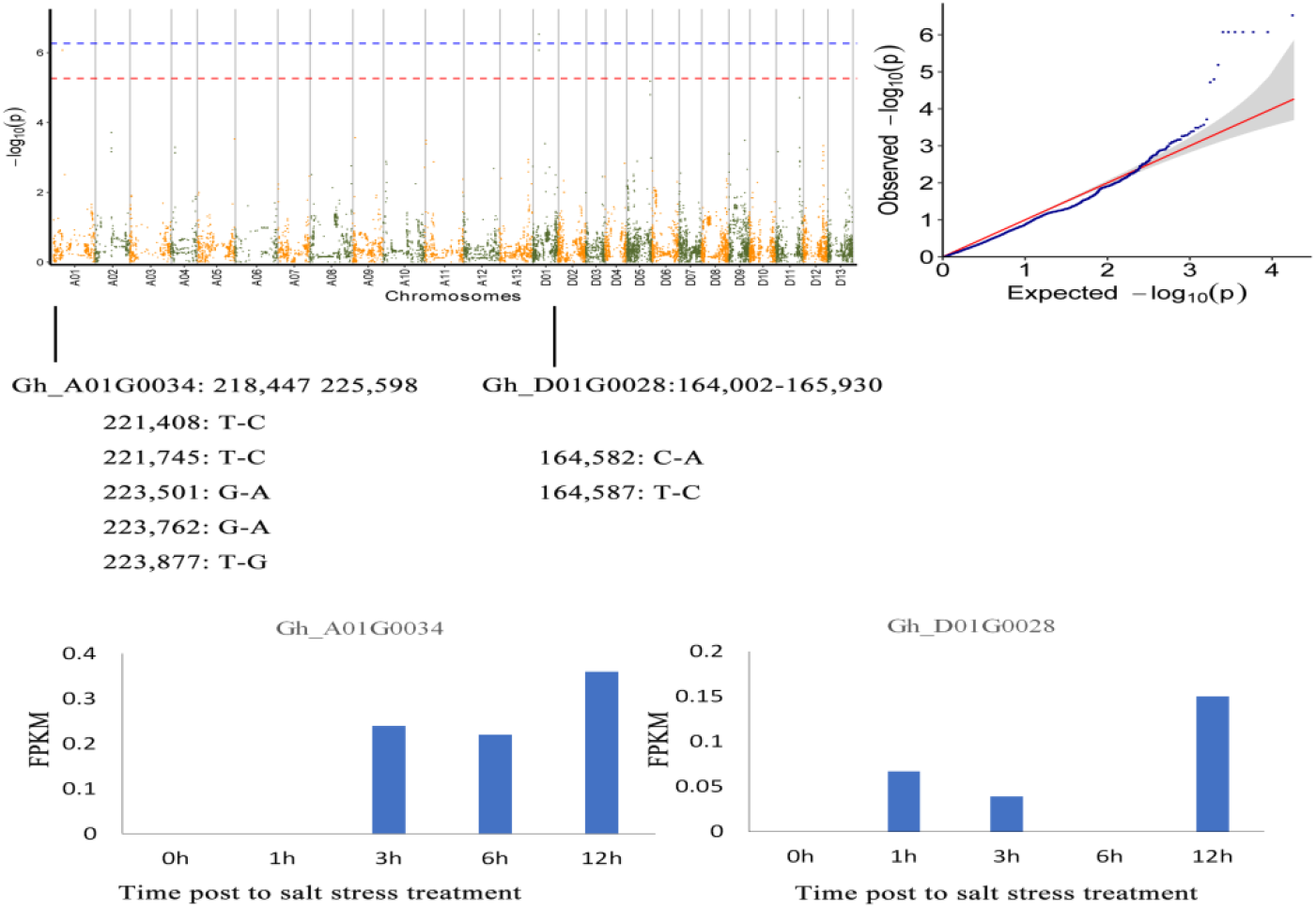
Screening of candidate genes for salt tolerance

### 2.6 Alignment of drought resistance-related genes and Arabidopsis homologous sequences

The comparison of Gh_A01G0034 with Tair3 and Arabidopsis showed that the homologous gene in Arabidopsis is AT4G34660, with 75% homology, and the gene is an SH3 domain-containing protein. The comparison of Gh_D01G0028 with Tair3 showed that the homologous gene in Arabidopsis thaliana is AT4G34940, with 73% homology, which encodes ARO1 armadillo repeat only 1 (Figure 8).

**Figure 8.**
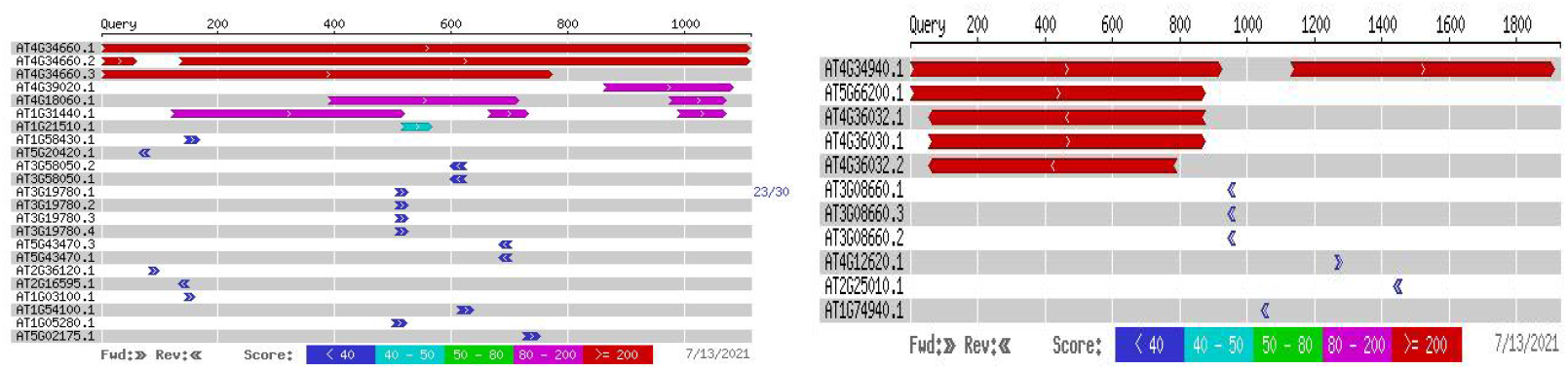
Gene sequence blast results

## 3 Discussion

### 3.1 Target gene identification based on GWAS

With the efficient development of genotyping technology, SNP markers have the advantages of wide distribution, high throughput, low cost and high accuracy. Genome-wide association analysis based on SNP genetic markers has become the first choice for analysing the complex traits of humans, animals and plants (Huang et al, 2014). Platt et al. believe that some significant SNP markers in linkage disequilibrium can show a higher degree of linkage disequilibrium than those SNPs that actually cause phenotypic variation (Platt et al, 2010, Korte et al, 2013).

Sun et al. (Sun et al, 2017) used the GWAS method to type SNPs in cotton crops using microarrays, developed 10,511 polymorphic SNP markers from a natural population composed of 719 materials, and associated 46 with significantly excellent sites related to fibre quality. Two haplotypes related to fibre length and fibre strength were identified on chromosomes At07 and Dtll. Reddy et al. used GBS (genotyping-by-sequencing) SNP typing technology to develop 10,129 polymorphic SNP markers from upland cotton and sea island cotton based on SNP markers and linkage disequilibrium LD from upland cotton and sea island cotton. A total of 142 and 282 blocks were excavated from sea island cotton (Reddy et al, 2017).

In this study, Illumina Cotton SNP 70K was used to develop 18,432 SNP markers in the whole genome. On this basis, whole-genome association analysis was used to associate excellent sites related to salt tolerance traits and the salt tolerance index.

### 3.2 Functional analysis of candidate genes

Candidate genes are a class of genes whose expression on the chromosome is not clear. They are involved in the phenotypic expression of organisms, and association analysis suggests that they are related to a certain part of the genome. Such genes may be structural genes, regulatory genes, or affect the expression of traits in biochemical metabolic pathways. The functional insufficiency of the candidate gene is known, and whether it is related to drought resistance has been verified. According to the screening, functional annotations can be assigned, or Arabidopsis homologous genes can be found from the gene information. This method has been previously reported to target genes that are clearly related to salt tolerance. GWAS analysis is a fast and powerful method to mine regulatory genes through crop indicators. In this study, two candidate genes related to salt tolerance, Gh_A01G0034 and Gh_D01G0028, and Arabidopsis homologous genes AT4G34660 and AT4G34940 were identified in the significant SNP sites.

Src homology-3 (SH3) domain-containing protein 2 (SH3P2), as a ubiquitin- and ESCRT-I-binding protein that functions in intracellular trafficking (Marie-Kristin, et al., 2017), plays a crucial role at the step of membrane tubulation during cell plate formation (Gyeongik Ahn, et al., 2017). SH3P2 colocalized with clathrin light chain- labelled punctate structures and interacted with the clathrin heavy chain in planta, indicating a role for SH3P2 in clathrin-mediated endocytosis (Marie-Kristin, et al., 2017). Clathrin-mediated endocytosis of plasma membrane proteins is an essential regulatory process that controls plasma membrane protein abundance and is therefore important for many signalling pathways, such as hormone signalling and biotic and abiotic stress responses (Li, 2020).

Armadillo repeat protein (ARO1) is one gene in a family of four in Arabidopsis. It is localized in the nucleus and cytoplasm of pollen vegetative cells and in the cytoplasm of egg cells and is involved in the signalling network, controlling tip growth and actin organization in the pollen tube. The signal-mediated and spatially controlled assembly and dynamics of actin are crucial for maintaining the shape, motility, and tip growth of eukaryotic cells. ARO1 is specifically expressed in the vegetative cells of pollen as well as in egg cells (Marina Gebert, et al, 2008).

## 4 Conclusion

A total of 18,432 polymorphic SNP markers were developed and screened from natural populations using gene chip technology. These SNP markers were used to analyse the structure of the population to obtain the Q matrix, and then the salt tolerance traits and salt tolerance index data were combined to conduct a genome-wide association analysis. The natural population can be divided into two subgroups. The genetic relationship between the materials was weak, indicating that the breed inherited diversity is decreasing. The salt tolerance traits were associated with 27 significant SNP sites, and the salt tolerance index was associated with 15 significant SNP sites. The significant SNP sites were further analysed, and the traits were associated with the 7 d shoot length decline rate. Salt tolerance-related alleles Gh_A01G0034 and Gh_D01G0028 were detected in the spot data. The transcriptome results showed that the expression of these two genes reached their peak at 12 hours of salt stress. The homologous sequences were compared with Arabidopsis thaliana to obtain the homologous genes AT4G34660 and AT4G34940. Analysis of the functions of these two genes revealed that the Arabidopsis thaliana homologous sequence encodes the SH3 domain protein and ARO1. The membrane lipid peroxidation scavenging system has high activity in the salt tolerance reaction of cotton, so the stability of the structure and function of the protective membrane is the key to the salt tolerance of cotton. This study further analysed the functions and expression patterns of cotton salt-tolerant genes and even has certain reference value for analysing the mechanism of cotton salt tolerance.

## Dota availability statements

The raw sequence data reported in this paper have been deposited in the Genome Sequence Archive (Genomics, Proteomics & Bioinformatics 2017) in National Genomics Data Center (Nucleic Acids Res 2021), China National Center for Bioinformation / Beijing Institute of Genomics, Chinese Academy of Sciences, under accession number CRAxxxxxx that are publicly accessible at https://ngdc.cncb.ac.cn/gsa.

## Acknowledge

We would like to thank Xinjiang Academy of Agricultural Sciences, China, for the cotton varieties provided for this study, BMK for the sequencing, and AJE for the English polishing. Thanks to all the people, units and enterprises who have provided help to this study.

## Contribution tribuauthor

Zeliang Zhang is the executor of the experimental design and experimental research of this study; Zeliang Zhang, Zhiwei Sang completed the data analysis, the writing of the paper; Junduo Wang, Yajun Liang, Quanjia Chen, Zhaolong Gong, Jiangping Guo participated in experimental design, experimental data collection and test results analysis; Xueyuan Li, Juyun Zheng is the architect and director of the project, guiding experimental design, data analysis, paper writing and modification. The entire authors read and agree to the final text.

**No competing interests among all authors.**

